# Effects of Liraglutide on Gut Bacterial Community Dynamics

**DOI:** 10.64898/2026.01.26.701836

**Authors:** Jami Bull, Paul L Durham, Babur S Mirza

## Abstract

Liraglutide, a GLP-1 receptor agonist, is used to induce weight loss. However, limited information exists on liraglutide’s effects on the gut bacterial community and their restoration after washout. We investigated liraglutide’s effect on the gut bacterial community in diet-induced obese (DIO) mice and whether these changes persist after washout. Twenty-four male C57BL/6J mice on high-fat (HFC) or low-fat (LFC) diets were monitored for 21 days. A subgroup of high-fat mice received daily liraglutide for 14 days (HFL), followed by a 7-day washout. Liraglutide induced significant weight loss by Day 4, which persisted during treatment and partially reversed post-treatment. For bacterial community analysis, 7.1 million 16S rRNA gene sequences were retrieved using Illumina paired-end sequencing. We observed distinct shifts in gut bacterial community structure during liraglutide treatment, which mostly returned to baseline after the 7-day washout. Using SIMPER analysis, 21 amplicon sequence variants (ASVs) were identified as major contributors. Nine ASVs, related to *Lactobacillus gasseri, L. paragasseri, L. johnsonii*, and *Leptogranulimonas caecicola*, significantly increased during treatment and declined post-washout. The remaining 12 ASVs, associated with protein- and carbohydrate-fermenting bacteria (*Romboutsia, Faecalicatena, Oscillibacter*), decreased during treatment. Comparison across all groups identified 29 ASVs, clustering into seven phylogenetic groups, highlighting liraglutide’s enrichment of bile-acid- and mucin-associated taxa and suppression of carbohydrate-fermentative genera. These findings demonstrate that liraglutide induces rapid, diet-dependent, yet reversible shifts in the gut microbiome, favoring lactic acid–producing bacteria while reducing fermentative taxa. Such microbial changes may contribute to liraglutide’s metabolic effects and provide insight into host– microbiome interactions in obesity treatment.

**IMPORTANCE:** Obesity and overweight states are intricately linked to the gut bacterial community, yet the effects of common obesity treatments such as GLP-1 receptor agonists on gut bacteria remain unclear. Here, we show that liraglutide, a GLP-1 analog, reshapes the gut bacterial community in diet-induced obese mice relative to untreated obese and lean controls. By including baseline samples when mice were at a normal weight, we distinguished bacterial changes due to the drug from those due to obesity progression, as well as how the community structure is affected during a washout period. Liraglutide treatment selectively increased beneficial gut bacteria (e.g., Lactobacillaceae) under high-fat conditions. These bacterial shifts during GLP-1 therapy may contribute to its metabolic benefits and broaden our understanding of host bacterial interactions in the context of diet and weight management.

## INTRODUCTION

The dramatic increase in the number of overweight and obese individuals over the past several decades poses global health concerns (1). While being overweight is defined as having a body mass index (BMI) over 25, obesity is associated with a value between 30-34.9. Their rising prevalence is associated with numerous comorbidities, including type 2 diabetes, prediabetes, hypertension, and cardiovascular disease (2). While these chronic conditions are primarily driven by a lack of physical activity and poor dietary choices, the rate of progression is influenced by a combination of genetic, epigenetic, and microbial factors (3–5). The complexity of these conditions highlights the need for a deeper understanding of the mechanisms contributing to their development, in particular, the imbalance in the gut microbiota that can alter metabolism, produce harmful metabolites, and promote inflammation.

With the rising rates of obesity, which is considered a chronic inflammatory disease, the development of therapeutic strategies to slow or reverse its progression is crucial. Towards this goal, drugs that mimic the two natural incretin gut hormones, glucagon-like peptide-1 (GLP-1) and glucose-dependent insulinotropic polypeptide (GIP), are approved by the FDA for use as anti-diabetic therapeutics and for weight loss. These drugs act as agonists at their receptors to stimulate insulin production and release, suppress glucagon secretion, slow gastric emptying, and suppress appetite (6). Liraglutide is an FDA-approved prescription medication for chronic weight management (7, 8). Similar to other incretin hormone drugs, chronic use is associated with an increased risk of negative side effects, including nausea, vomiting, diarrhea, abdominal pain, and constipation (9, 10). So far, limited information is available on how liraglutide regulates gut microbial imbalance (dysbiosis), which has been linked to the development and persistence of obesity and other related diseases (11). Although dysbiosis is not considered a direct cause of obesity, evidence shows distinct shifts in the gut bacterial composition of obese individuals, characterized by lower bacterial diversity and reduced gene richness compared to lean individuals (12). These shifts indicate a complex relationship between gut bacterial composition and metabolic health.

The gut bacterial community plays a significant role in regulating host metabolism, energy balance, and body weight (13) with beneficial commensal bacteria supporting immune function, gut barrier integrity, and metabolic homeostasis (14, 15). A decrease in beneficial bacteria may contribute to increased oxidative stress and metabolic disturbances (16) with an elevated F/B ratio being reported in obese mice compared with their lean counterparts (17). However, this ratio is not always consistent and may vary based on dietary composition, such as purified high-sugar diets that can independently alter microbial composition (18).

Animal models, such as diet-induced obese (DIO) mice, have been crucial in studying the role of the gut bacterial communities in obesity and other comorbid conditions. DIO mice serve as a relevant preclinical model for studying obesity in humans, which is promoted primarily by dietary factors rather than genetic mutations. Data from preclinical models have identified the presence of obesity-associated bacterial species (19) that can alter the ability to extract energy and nutrients from the diet, regulate host metabolism, and body weight (4, 13). In this study, changes in the gut microbiota were investigated in male mice maintained on high-fat or low-fat diets that received daily injections of liraglutide. Bacterial community analysis of high-quality 16S rRNA gene sequences generated from Illumina paired-end sequencing was performed to identify the key bacterial genera that were altered in response to liraglutide treatment. Our findings provide evidence that liraglutide induces rapid, diet-dependent, yet reversible shifts in the gut microbiome, favoring lactic acid-producing bacteria while reducing fermentative taxa.

## RESULTS AND DISCUSSION

### Liraglutide induced changes in mice weight over time

Similar body weights were observed among all animals at the start of liraglutide treatment across all treatment groups (**Fig. 1; Table S1**). Based on the standardized Δ weight analysis (mean weight difference), mice in the high-fat control (HFC) group initially maintained their body weight for a few days and then showed a significant increase by Day 7, which continued throughout the remainder of the study (**Fig. 1**). In contrast, mice in the high-fat liraglutide (HFL) treated group experienced a reduction in body weight, and by Day 4 of liraglutide treatment their Δ weight values were significantly lower compared to the HFC group. This difference remained significant for the remainder of the study (**Fig. 1**). Overall, the HFL group showed a clear decrease in Δ weight over the first 14 days of daily liraglutide treatment and remained significantly lower compared to the HFC group. After stopping liraglutide treatment on Day 14, a gradual increase in weight from Day 17 through Day 21 was observed. This finding demonstrates that liraglutide’s weight-reducing effect was transient. The LFC group maintained relatively stable body weight and Δ weight values throughout the study, with a significant increase observed toward the end (**Fig. 1**).

**Fig. 1.**
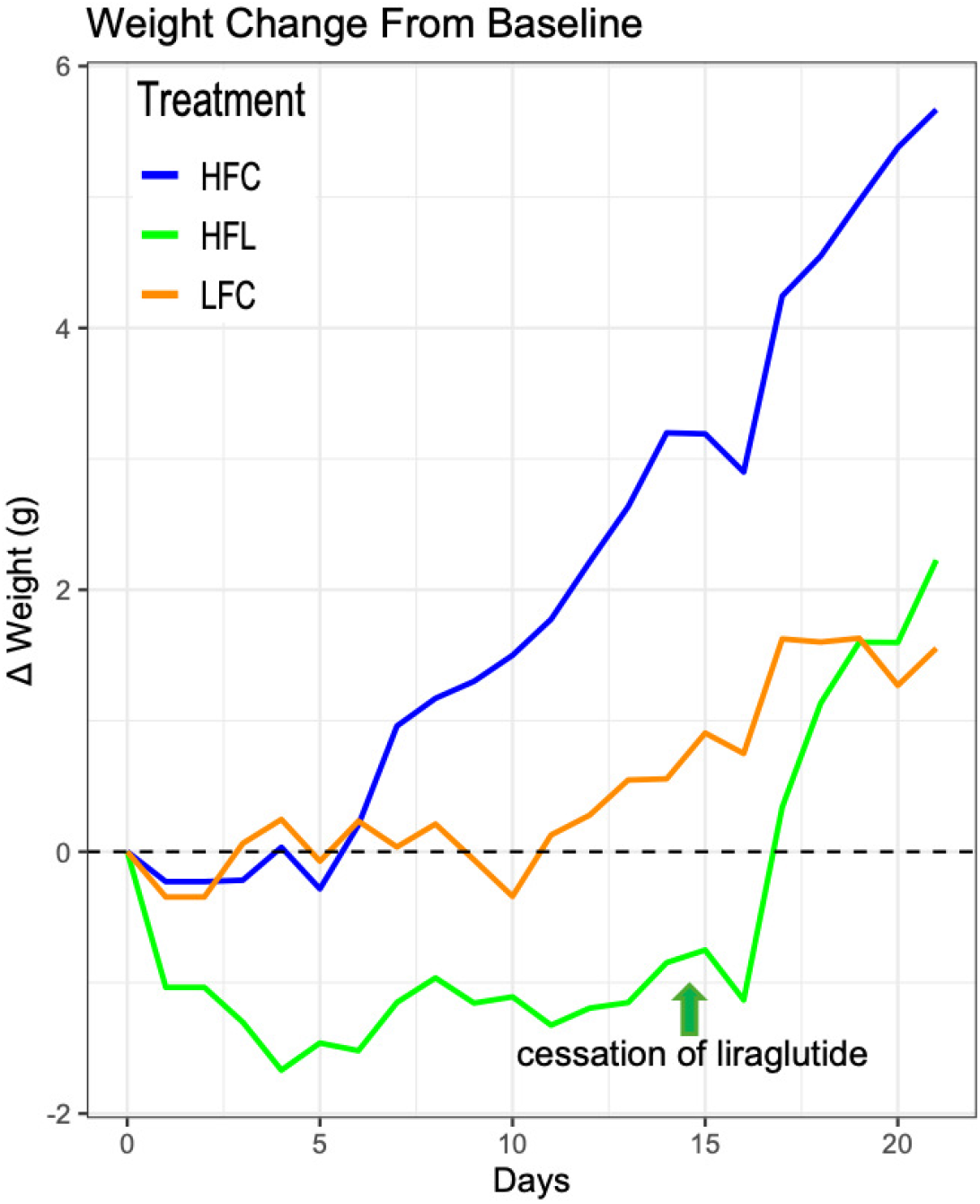
Body weight change was calculated by subtracting each group’s Day 0 mean body weight from the corresponding daily mean values. Data are presented as the means for each treatment group. HFL (liraglutide-treated) mice exhibited reduced Δ weight values throughout the treatment period, whereas HFC mice showed normal progressive weight gain. Following cessation of liraglutide on Day 14, a regaining of body weight was observed in the HFL group.

Similar to our results, previous studies (20–22) also demonstrated liraglutide’s role in promoting weight loss in mice. Liraglutide primarily acts as a GLP-1 receptor agonist, mimicking the effects of the naturally occurring hormone GLP-1. Some of the suggested mechanisms of weight loss include suppression of appetite, delay gastric emptying, and increased satiety (23, 7, 24). The weight loss was not sustainable as an increase in weight was observed during the washout period (**Fig. 1**).

### Phylum level distribution of bacterial sequences

A total of 7.1 million high-quality bacterial 16S rRNA gene sequences were classified. Most of the sequences (6,297,930 sequences, accounting for 88.7% of the total) were related to Firmicutes, followed by Bacteroidetes (7.7%) and Actinobacteria (1.8%) (**Fig. 2**). Sequences related to other bacterial phyla were retrieved at lower abundances (<1% of the total sequences; **Fig. 2**). Firmicutes are commonly associated with the fecal microbiota of warm-blooded animals including humans (25). In mice, elevated Firmicutes levels are often linked to obesity, although many species within this phylum contribute positively to gut homeostasis (26, 27). Bacteroidetes were the second most abundant phylum, comprising 7.7% of sequences (115,389 sequences) (**Fig. 2**). Members of Bacteroidetes are frequently associated with metabolic health and leanness; however, certain genera within this phylum may not be beneficial (26, 28).

**Fig. 2.**
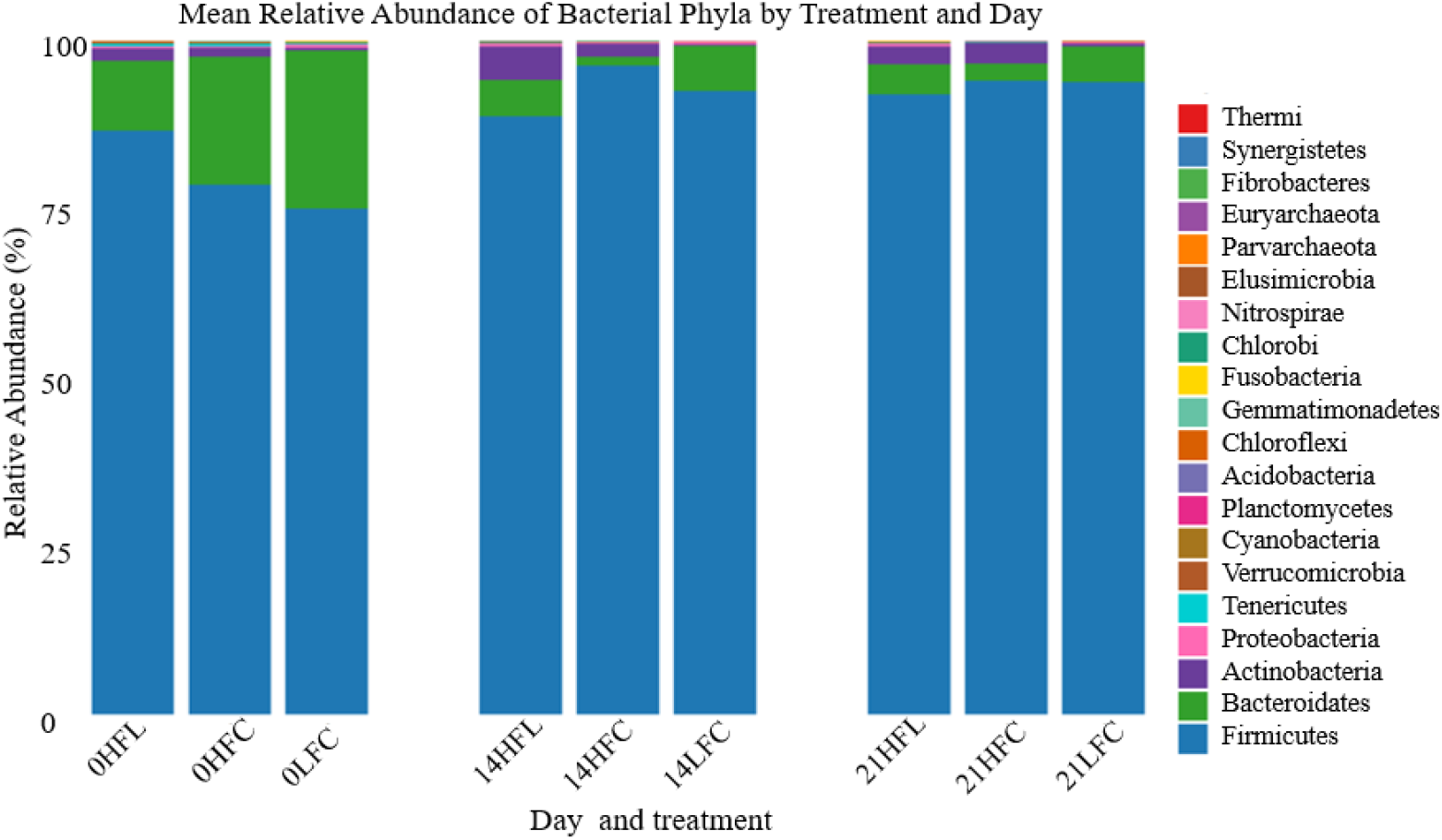
Phylum-level distribution of bacterial 16S rRNA gene sequences across three treatment groups (HFL, HFC, LFC) at three time points (Days 0, 14, and 21).

### Liraglutide induced shifts in gut bacterial community structure

Considering the decrease in animal weight in response to liraglutide treatment, one of the main questions of our study was to assess if bacterial community structure is altered during liraglutide exposure and whether the bacterial community could be restored after ceasing liraglutide treatment. Overall, approximately 7.1 million high-quality 16S rRNA gene sequences across all treatments (**Table S1**) were retrieved and significant changes in bacterial community structure were observed based on non-metric multidimensional scaling (NMDS) analysis in response to both liraglutide treatment and diet (**Fig. 3A–C**). Specifically, a distinct shift in the bacterial community structure in HFL treatment group was observed after 14 days of liraglutide treatment when compared to both its baseline community at Day 0 and the community observed at Day 21 following the 7-day washout period (**Fig. 3A**). Likewise, the bacterial community of the HFL group at Day 14 was significantly different from both the HFC and LFC groups (**Fig. 3B**). Finally, by Day 21 (after the 7-day washout period), the bacterial community of the HFL group resembled that of the HFC group but remained significantly different from the LFC group, suggesting dietary influence on the gut bacterial community (**Fig. 3C**).

**Fig. 3.**
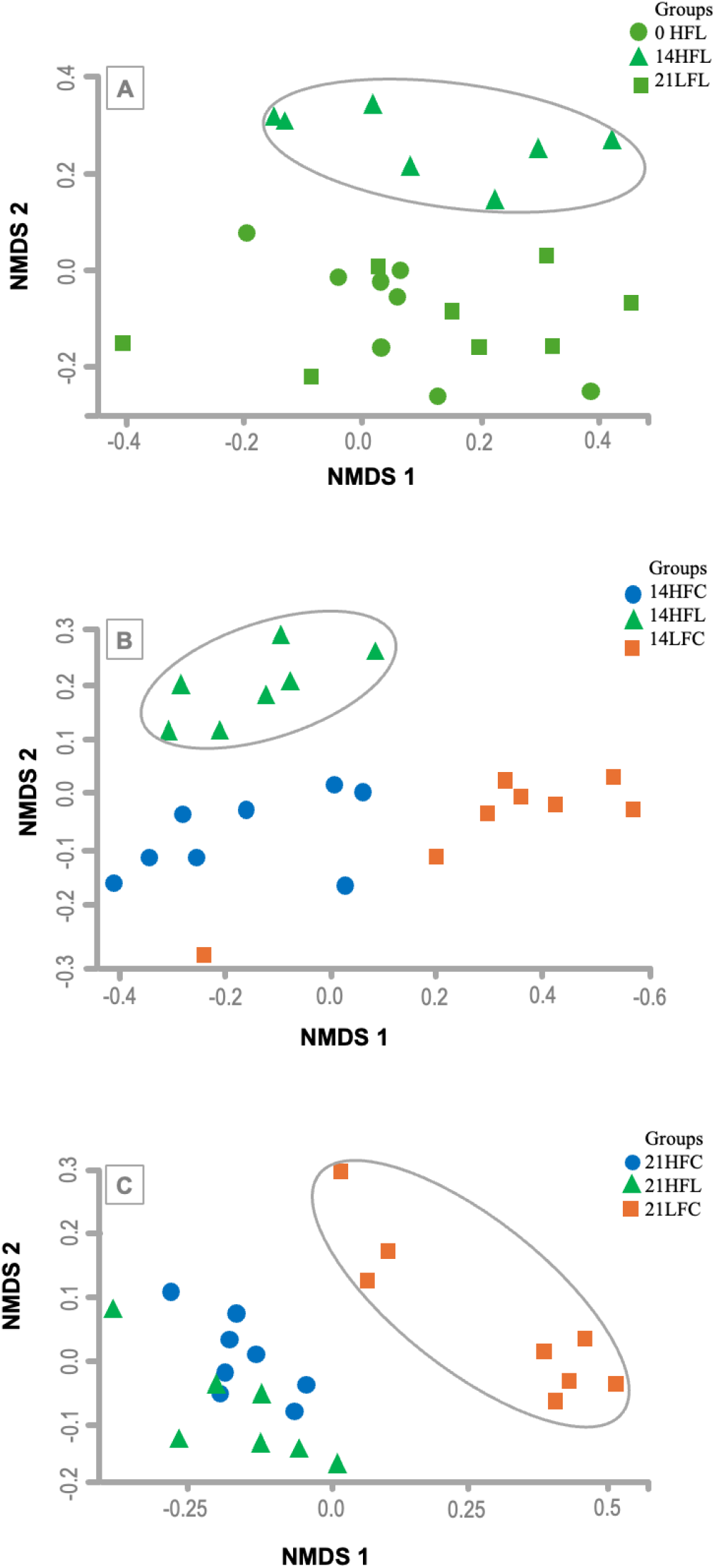
**A–C.** Non-metric multidimensional scaling (NMDS) plots based on Bray–Curtis similarity of 16S rRNA gene sequences (97% similarity). Panel **(A)** shows the NMDS ordination of the HFL group across all time points (Day 0, Day 14, Day 21), highlighting the transient shift in community composition during liraglutide treatment and its return toward baseline following washout. Panel **(B)** presents the NMDS ordination of all three treatment groups on Day 14, illustrating clear differences in bacterial community structure during liraglutide exposure. Panel **(C)** shows the NMDS ordination of all three treatment groups on Day 21, demonstrating the post-treatment convergence of the HFL group with the HFC group following the washout period. Each point represents an individual sample, and distances between points reflect the degree of compositional dissimilarity among bacterial communities. HFL (high-fat diet + liraglutide), HFC (high-fat control), LFC (low-fat control).

This shift in the bacterial community in response to liraglutide could be due to several previously suggested indirect mechanisms (**Fig. 3A**), such as liraglutide altering the gastrointestinal environment by slowing gastric emptying (29), reducing food intake by promoting satiety (30), and altering the release of bile acids and pancreatic enzymes (31). Additionally, liraglutide has also been suggested to exert direct effects on the bacterial community by limiting substrate availability (32, 33) for bacterial fermentation and by altering gut epithelial function and mucosal immunity (34).

Unlike previous studies, we also assessed the bacterial community following a 7-day washout period without liraglutide and observed a restoration of the bacterial community in the HFL group (Day 21), which closely resembled the original community at Day 0 (**Fig. 3A**), and that of the HFC group across different time points (Day 21 comparison is shown in **Fig. 3C**). This restoration suggests that liraglutide-induced shifts in the gut bacterial community structure are transient and not maintained in the absence of treatment. Although the precise mechanisms by which liraglutide alters the gut microbial community remain unclear, the reversibility of the gut microbiome after cessation of treatment suggests a strong treatment-dependent influence on the animal gut bacterial community. Lastly, as anticipated, we observed significant differences in bacterial community structure between the HFC and LFC groups, indicating a strong dietary influence on the bacterial community in the mouse gut.

### Identification of bacterial species altered in responses to liraglutide treatment

As discussed above, the NMDS analysis clearly demonstrated a distinct shift in the bacterial community of the HFL group on Day 14, followed by a restoration toward the original bacterial community by Day 21, similar to that observed at baseline (Day 0). Another important question addressed in our study to identify the specific bacterial species that were altered in response to liraglutide treatment.

Hence, a SIMPER analysis was conducted that identified 21 ASVs, which contributed approximately 83% of the variation observed in bacterial community structure in response to liraglutide treatment within the HFL group across different time points. Nine of these 21 ASVs (representing 1,274,508 sequences; 64% of total sequences in the HFL group) showed a significant increase in abundance during liraglutide treatment and returned to baseline levels by Day 21 following the 7-day washout period (**Fig. 4**).

**Fig. 4.**
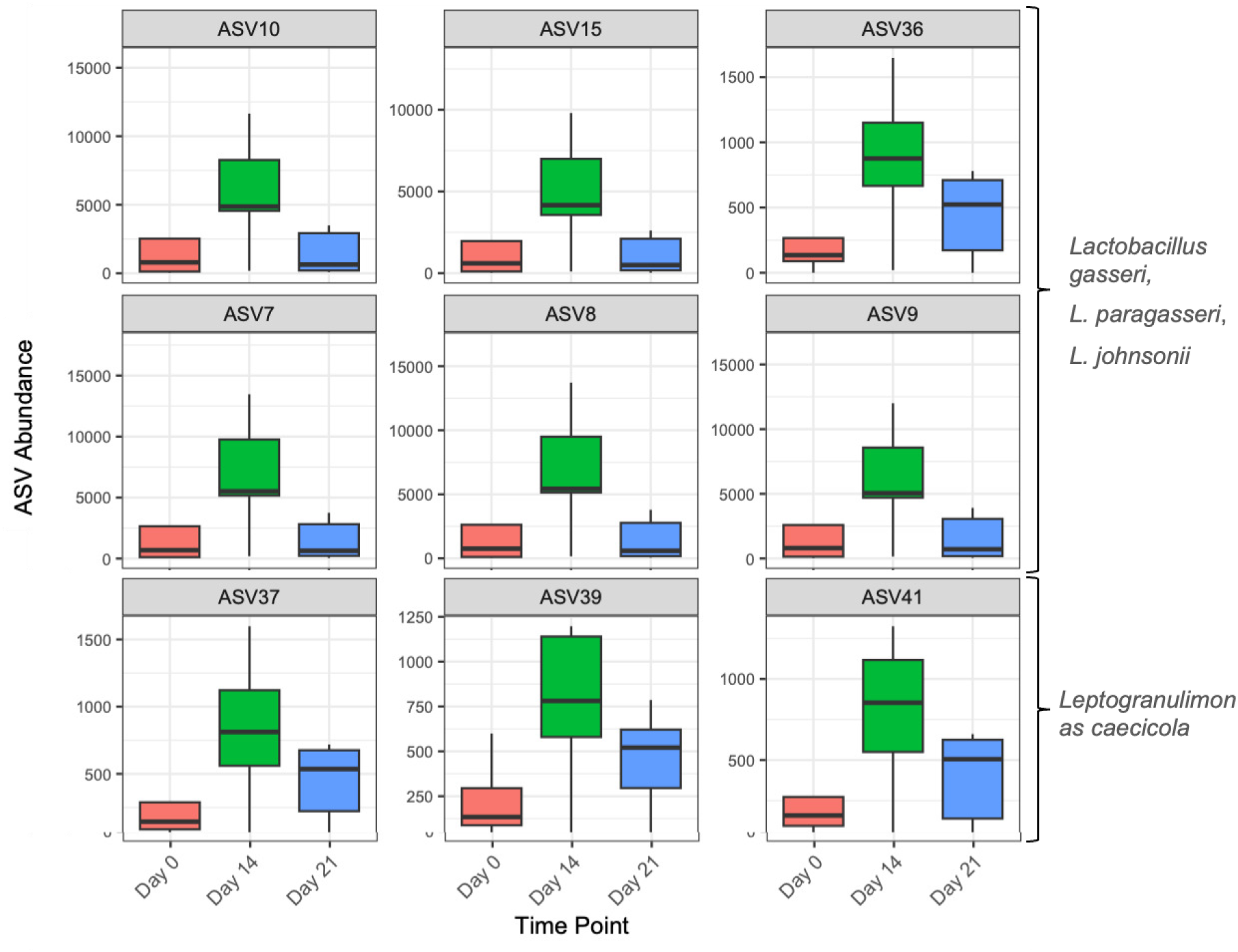
Boxplots showing the average sequence abundance of nine ASVs that significantly increased in response to 14 days of liraglutide treatment and subsequently decreased after a 7-day washout period. The six ASVs in the upper two panels are related to *Lactobacillus* species, while the three ASVs in the lower panel are associated with *Leptogranulimonas caecicola*. Each box represents the distribution of sequence counts across eight fecal samples at each time point.

Furthermore, to determine whether the changes in these nine ASVs in the HFL group were age-related or induced by liraglutide, the relative distribution of the same nine ASVs in the HFC group across the same time points (Days 0, 14, and 21) was assessed. No significant trend of an increase on Day 14 followed by a decrease on Day 21 was observed in these ASVs in the HFC group (**Fig. S2**), which further supports the notion that liraglutide can play a direct or indirect role in influencing the gut microbiome. Additionally, a much lower relative abundance (10.2% of total sequences in the HFC group) of sequences corresponding to these nine ASVs in the HFC group was observed when compared to the HFL group, while the relative abundance of sequences related to these nine ASVs was very low (<1%) in the LFC group.

Through NCBI blast search of representative sequences from the nine ASVs, six of the nine identified ASVs (ASV 7-10, 15, 36) were affiliated with (∼99% sequence similarity) closely related *Lactobacillus* spp. (*L. gasseri, L. paragasseri, and L. johnsonii*) (**Fig. 4**; **Table S2**). *Lactobacillus* spp. such as *L. gasseri* and *L. johnsonii* are well-established beneficial bacteria reported to support host health through multiple mechanisms such as production of lactic acid, mucosal adhesion, pathogen exclusion by acid production, and also development of host immune system (35–41).

*Lactobacillus* spp. also ferments complex carbohydrates to yield short-chain fatty acids (SCFA), which function to maintain gut barrier integrity, regulate immune responses, influence host metabolism, and gut-brain signaling (42). The transient rise of *L. gasseri* in response to liraglutide may be mediated by delayed gastric emptying and enhanced mucin secretion, both of which favor fermentative, mucin-associated bacteria (29,34, 43).

The remaining three of the nine ASVs (ASV 37, 39, 41) were closely related to *Leptogranulimonas caecicola*. These ASVs showed a slight increase in the HFL group at Day 14 (3.9%) compared with baseline (1.3%) and controls (1.4%). *L. caecicola* is also known for lactic acid production and possesses bile salt hydrolase enzyme activity (44). The observed *L. caecicola* enrichment was likely due to liraglutide-driven changes in bile acid composition and intestinal pH.

Additionally, 12 ASVs (containing 4.41% of total sequences in the HFL group across the three time points; 88,361 sequences) were identified that showed a significant decrease in abundance in response to liraglutide treatment on Day 14 compared with baseline (HFL) and with both HFC and LFC treatments (**Fig. S**2). Overall, the sequences within these 12 ASVs were less abundant (4.41%) compared with the 64% of total sequences in the HFL group that were related to the 9 ASVs which showed an increase in response to liraglutide. Specifically, within the HFL treatment group, these twelve ASVs represented 11.49% of all sequences in HFL at baseline (Day 0) but declined sharply to 2.61% of all HFL sequences on Day 14, with partial recovery to 8.54% by Day 21 following the 7-day washout period. In contrast, this strong suppression pattern was not observed in either control group. In the HFC mice, the same twelve ASVs showed no consistent decrease over time, representing 9.47%, 4.30%, and 5.30% of total sequences at Days 0, 14, and 21, respectively. The LFC group exhibited consistently low relative abundances (<3%) across all time points, with abundances falling below 0.4% during Days 14 and 21. Together, these findings indicate that liraglutide specifically suppresses these bacterial species during treatment, and that this suppression is reversible once the treatment is discontinued.

The sequences within these 12 ASVs were related to *Lactococcus lactis, Romboutsia ilealis, Faecalicatena fissicatena, Clostridiales* bacterium CIEAF 017, and *Oscillibacter* sp. Members of *Romboutsia, Faecalicatena*, and *Oscillibacter* are commonly associated with protein and amino-acid fermentation and often expand under high-fat dietary conditions (45– 49). Their suppression during liraglutide treatment therefore likely reflects liraglutide-induced shifts in host energy intake and altered substrate availability. Their recovery after liraglutide treatment withdrawal further supports a strong influence of liraglutide on the gut bacterial community.

### Phylogenetic clustering or comparison of gut microbiome in responses to all three treatments (HFL, HFC and LFC)

Furthermore, the key ASVs that mainly contributed toward the overall observed variation across three different treatments and time points were assessed. ASVs were compared through pairwise SIMPR analysis within different treatments, and 29 ASVs were identified that contributed to ∼80% of observed variation across the three treatment groups.

Representative sequences from these ASVs along with the related sequences from NCBI were phylogenetically compared and grouped into seven major clusters (**Fig. 5**). All sequences were from phylum Firmicutes and closely related to species such as *Lactobacillus* spp., *Faecalibaculum rodentium, Dubosiella newyorkensis, Romboutsia timonensis, Acetatifactor muris, Leptogranulimonas caecicola*, and *Duncaniella muris* (**Fig. 5**).

**Fig. 5.**
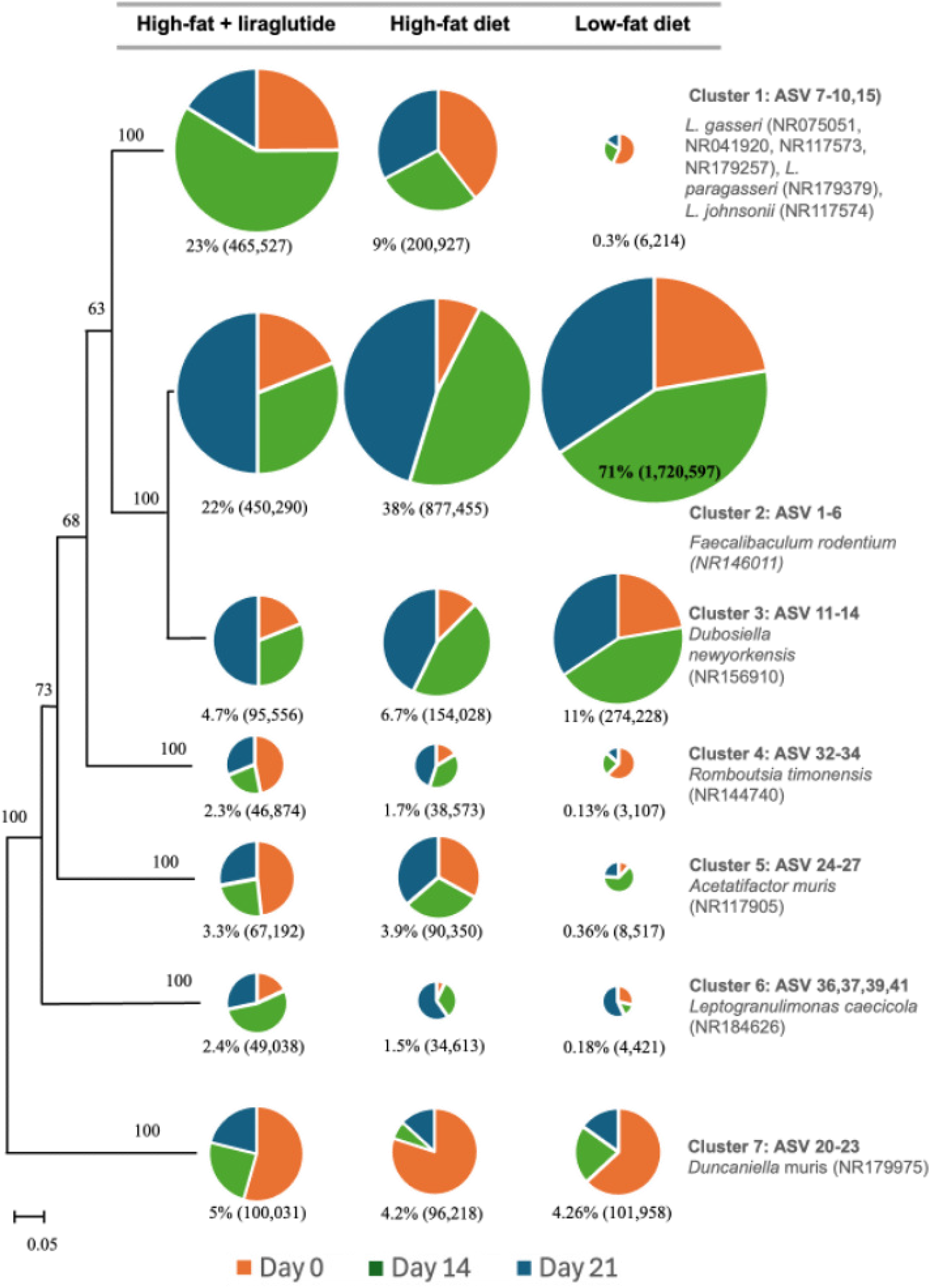
Maximum likelihood phylogenetic tree based on 16S rRNA gene sequences representing 29 ASVs that together contributed nearly 80% of the observed variation across treatments and sampling time points. Samples were collected from each treatment group (HFL, HFC, LFC) at three times: Day 0 (tan), Day 14 (green), and Day 21 (blue). Only ASVs representing ≥10,000 sequences and contributing significantly to differences among treatments and time points are included. The size of each pie chart is proportional to the total number of sequences within that ASV cluster, and the percentage contribution of sequences from each treatment group is shown beneath each pie. Overall, sequences were grouped into seven clusters based on high nucleotide similarity and strong bootstrap support. Bootstrap values (≥50%) are shown at corresponding nodes, and branches within clusters were collapsed for clarity. Closely related reference sequences from NCBI included for comparison are listed to the right of the phylogenetic tree.

As described above, in the HFL group, sequences related to *Lactobacillus* spp. and *Leptogranulimonas caecicola* were grouped into clusters 1 and 6, respectively, and both showed significant increases in response to liraglutide treatment (**Fig. 4**). Specifically, at baseline (Day 0), *Lactobacillus* spp. accounted for 17.1% of total sequences in the HFL group, which increased markedly to 41.0% on Day 14. This enrichment declined to 11.5% by Day 21 (washout). In contrast, *Lactobacillus*-related sequences represented 9.0% of total sequences in the HFC group at baseline, decreasing to 6.7% and 6.9% on Days 14 and 21, respectively (**Fig. 5**). In the LFC group, *Lactobacillus*-related sequences remained very low (< 1%) throughout the experimental period. Likewise, *Leptogranulimonas caecicola* (cluster 6) showed a modest increase in liraglutide-treated mice (HFL) at Day 14 (3.9%) compared to baseline (1.3%) and high-fat controls (1.4%). As discussed above, both *Lactobacillus* spp. and *L. caecicola* produce lactic acid and expresses bile salt hydrolase, suggesting a role in bile acid metabolism and intestinal pH regulation (44).

In contrast to the *Lactobacillus* and *L. caecicola* enrichment observed with liraglutide treatment, *Faecalibaculum rodentium* (cluster 2) was detected across all dietary groups but detected in higher abundance in the LFC group mice, comprising 70% and 60% of total sequences within LFC treatment at Days 14 and 21, respectively (**Fig. 5**). Although abundant in both control diets, its markedly lower representation in the liraglutide-treated high-fat group (HFL) suggests that liraglutide may indirectly suppress carbohydrate-fermenting taxa by reducing nutrient intake and altering substrate availability (33, 50). *F. rodentium* is a fermenting bacterium that produces SCFA commonly associated with carbohydrate metabolism and energy harvest in mice (51, 52). The reduced abundance of this genus during liraglutide treatment may therefore reflect a shift toward a less fermentative gut environment, due to less food uptake and delayed gastric emptying frequently associated with liraglutide treatment (29).

Similarly, *Dubosiella newyorkensis* (cluster 3) increased notably in LFC mice (11% sequences within LFC group) but was less prevalent in liraglutide-treated groups (4.7% of sequences in HFL group). *D. newyorkensis* has been suggested as a potential probiotic against age-related diseases (53) and may play a role in immune tolerance in colitis problems (54). Conversely, the relative abundances of *Acetatifactor muris* and *Duncaniella muris* (cluster 5 and 7) declined following liraglutide exposure. Both genera are members of the *Lachnospiraceae* family and are suggested to contribute to butyrate and acetate production (55, 56). Their reduction in the HFL group may indicate altered fermentation dynamics or a modification of SCFA production pathways in response to liraglutide-induced changes in gut physiology.

Collectively, these compositional shifts suggest that liraglutide selectively favors bile acid and mucus membrane associated bacteria such as *Lactobacillus* and *Leptogranulimonas*, while suppressing carbohydrate-fermenting genera including *Faecalibaculum, Dubosiella, Acetatifactor*, and *Duncaniella*. This pattern is consistent with a transition toward a gut environment characterized by reduced carbohydrate fermentation and enhanced mucosal metabolism that may contribute to liraglutide’s metabolic and anti-inflammatory benefits (34, 43).

## MATERIALS AND METHODS

### Experimental Design and Mouse Treatment

In the current study, 24 male C57BL/6J diet-induced obese (DIO) mice were obtained from The Jackson Laboratory (Bar Harbor, ME) mice. Female mice were excluded because DIO females do not consistently gain weight under high-fat diet conditions, and the Jackson Laboratory does not offer female DIO models. Sixteen mice were maintained on a high-fat diet (60% kcal from fat, D12492; Research Diets) and are hereafter referred to as the high-fat control group (HFC). Eight mice were maintained on a low-fat control diet (LFC; 10% kcal from fat, D12450B; Research Diets) and are referred to as the low-fat control group. The Jackson Laboratory placed the mice on their respective diets starting at six weeks of age, and mice were purchased at 10 weeks of age. The mice were continued on their respective diets, as initiated by the Jackson Laboratory, for the entire duration of the study. All mice had *ad libitum* access to food and water. Food was weighed daily; however, the high-fat DIO mice frequently shredded their food, so food intake data were not further analyzed in this study.

Upon arrival, mice were acclimated for 14 days in the Missouri State University vivarium. Eight of the 16 high-fat diet (HFL) mice received a daily subcutaneous injection of liraglutide (0.2 mg/kg dissolved in phosphate buffered saline (PBS)) for 14 days. The remaining 8 low-fat diet and 8 high-fat diet mice received daily PBS injections. Fecal samples were collected at three time points: prior to treatment, following the 14-day drug administration period, and after a 7-day washout period (**Fig. S3**). Body weight was recorded daily throughout the study. A 7-day washout period was included to assess changes in gut bacterial composition after discontinuation of liraglutide. All animals were housed in the Missouri State University vivarium and maintained according to standard operating procedures established for the facility and mouse species. All procedures used in this study were reviewed and approved by the Institutional Animal Care and Use Committee (Protocol # 2024-18).

### Data Collection and PCR

Fresh fecal samples were collected from all mice during weighing, as they defecated instinctively while being handled. DNA was extracted using DNeasy PowerLyzer PowerSoil Kit (Qiagen, Germantown, Maryland), following the manufacturer’s instructions. The PCR amplification and DNA processing were performed as previously described (57). Briefly, the bacterial 16S rRNA gene was amplified using universal primers 515F and 907R, which target the bacterial 16S rRNA gene. Each 25 µL PCR reaction contained 1X buffer, 0.2 µM of each primer, 2.0 mM MgSO_4_, 0.2 µM dNTPs, 1.0 µL template DNA, and 5 units of Platinum High-Fidelity Taq polymerase (Thermo Fisher Scientific, Waltham, MA, USA). The first PCR cycle began with an initial denaturation at 95°C for 5 minutes, followed by 30 cycles of 95°C for 45 seconds, annealing at 56°C for 45 seconds, extension at 72°C for 45 seconds, and a final extension at 72°C for 7 minutes. Positive (*E. coli* DNA) and negative (PCR-grade water) controls were run alongside the samples. PCR sample amplification was verified via gel electrophoresis and visualized using ethidium bromide staining.

Amplified products were cleaned using the ExoSap-IT PCR Cleanup System (Thermo Fisher Scientific), following the manufacturer’s protocol, and used as templates for the second PCR. In the second PCR, Illumina sequencing adapters A and B, along with unique multiplex identifiers, were incorporated. The PCR conditions were as follows; 95°C for 3 minutes, followed by 10 cycles of 94°C for 30 seconds, annealing at 60°C for 30 seconds, and extension at 72°C for 30 seconds, with a final extension at 72°C for 7 minutes. Amplified products were quantified using a NanoDrop 2000 spectrophotometer, pooled in equimolar concentrations, purified with Agencourt AMPure beads (Beckman Coulter, Brea, CA, USA), and sequenced using Illumina MiSeq paired-end sequencing. DNA quality and concentration were assessed using NanoDrop 2000.

### Data Analysis

Raw sequences were processed for quality control, including filtering by read length filtering (370-375 bp), removal of sequences containing more than eight ambiguous bases (N > 8), and removal of chimeric sequences using QIIME 2.0. Changes in bacterial community structure in response to treatment were analyzed at various taxonomic levels between treated and untreated control mice using USEARCH and RStudio. Quantification of bacterial 16S rRNA gene sequences enabled the identification of bacterial phyla and genera present at each time point, providing insight into potential dysbiosis.

### Statistical Analysis

We collected daily body weight and standardized weight trajectories across animals and treatment groups. Weight change from baseline (Δ weight) was calculated for each mouse relative to its Day 0 body weight. For each treatment group, the mean Δ weight for each day was computed by subtracting the Day 0 mean weight from the corresponding daily mean weight value. Δ weight values were then used to generate longitudinal weight-change curves to compare treatment-dependent growth patterns and to evaluate the effect of liraglutide (HFL) relative to the HFC and LFC control groups. Likewise, standard errors for Δ weight were derived from the standard errors of the raw weight measurements. The statistical significance of differences in body weight across treatments was assessed through repeated-measures ANOVA using R (58).

Overall bacterial community structure was assessed using the Bray-Curtis dissimilarity matrix. To evaluate the significance of differences in bacterial community structure across different time points, an analysis of similarity (ANOSIM) was conducted. Non-metric multidimensional scaling (NMDS) analysis was performed in RStudio 4.4.1 (58) using the vegan package (for ecological statistics), readxl (for importing Excel data), and ggplot2 (for visualization). The Bray-Curtis dissimilarity matrix was generated using the vegdist() function, and NMDS was performed using metaMDS() from the vegan package.

To identify the key ASVs contributing to major differences in the bacterial community after 14 days of liraglutide treatment, we performed a Similarity Percentages (SIMPER) analysis using R. This analysis identified 21 ASVs that accounted for most of the variation (>80%) across the different time points in the HFL treatment. Of these, nine ASVs significantly increased, and twelve ASVs significantly decreased in response to liraglutide treatment on Day 14. These differentially detected ASVs were then visualized as boxplots in R to illustrate their relative sequence abundance across time points.

Furthermore, samples across the other two treatments (HFC and LFC) were compared and overall 29 major ASVs were identified that contributed to the observed differences across all three treatments and sampling periods. Finally, a Maximum likelihood phylogenetic tree was constructed of representative ASV sequences and closely related sequences from GenBank using Molecular Evolutionary Genetic Analysis (MEGA 7) (59). Phylogenetic trees were generated with 1,000 bootstrap replicates using the Tamura-Nei model. Sequences showing high similarity (>98%) and strong bootstrap support were collapsed and grouped into seven major clusters (cluster 1 to 7). We also displayed the relative abundance of sequences within each cluster across treatments and time points using associated pie charts.

### Nucleotide sequence accession numbers

The 16S rRNA gene sequences were deposited in the NCBI Sequence Read Archive (SRA) under Bio project: PRJNA1344306

## ACKNOWLEGEMENTS

Thank you to Angela Goerndt and Jade Morris of the Missouri State University Vivarium for their expert guidance and technical support throughout this project.

Additional appreciation is extended to Irum Qureshi for her assistance with DNA extractions and sequencing library preparation.

